# An information content principle explains regulatory patterns of gene expression across human tissues

**DOI:** 10.64898/2026.02.19.706555

**Authors:** Ruthie Golomb, Maayan Yoles, Simon Fishilevich, Bar Cohen, Sapir Savariego Peled, Dvir Dahary, David Gokhman, Yitzhak Pilpel

## Abstract

Gene expression patterns range from broadly expressed housekeeping genes to highly tissue-specific ones. Notably, many genes exhibit intermediate specificity, characterized by elevated expression in some tissues while low or absent in others. Understanding how regulatory demands scale with tissue specificity offers a valuable opportunity to uncover fundamental principles of genome regulation. By analyzing *cis*-regulatory element (CRE) counts across human genes with varying tissue specificity, we observed a nonlinear pattern: genes with intermediate specificity harbor the highest CRE count, suggesting distinct regulatory strategies across the expression spectrum. Motivated by this observation, we used the Minimum Description Length (MDL) principle from information theory, together with a maximum parsimony approach from phylogenetics, to quantify regulatory demands across tissues.

Our analysis revealed that MDL-based regulatory demand scales consistently with diverse regulatory features, including CRE count, transcription-factor and microRNA targeting, and gene structure. To test whether this scaling changes across the expression spectrum, we partitioned genes by expression breadth. Two patterns emerged: features scaling with MDL in selectively expressed genes tend to act as on/off switches, whereas those in ubiquitous genes serve as fine-tuning knobs. Evolutionary analysis revealed that these regulatory patterns vary with gene age, with alignment between MDL and CRE counts peaking in intermediate-aged genes. Collectively, these results establish MDL combined with maximum parsimony as a powerful framework linking regulatory architecture, expression specificity, and evolutionary age, offering novel insights into the organizational principles underlying genome regulation.

## Introduction

Gene expression patterns vary widely across human tissues, reflecting distinct functional demands across cell types. Some genes are ubiquitously expressed, performing essential housekeeping roles such as RNA processing, protein synthesis, and core metabolism^1,2^. Others exhibit tissue-specific expression associated with specialized functions like neuronal signaling, spermatogenesis, and sensory perception^1,3^. Yet, between these extremes lies a large class of genes with intermediate tissue specificity, expressed highly in some tissues, yet minimally or not at all in others. Intermediately expressed genes have received less systematic attention, and their biological roles remain less characterized. Notably, genome-wide profiling by Yanai et al. demonstrated that this ‘midrange’ class comprises a substantial fraction of genes and suggested that crucial functional information resides within them^4^. Despite their abundance, it is unclear whether the characteristics of intermediately expressed genes simply represent a midpoint between broadly and narrowly expressed genes, or whether they exhibit distinct, even exaggerated, features.

Patterns of gene expression are governed by multiple interconnected layers of regulation. Central to this regulatory architecture are cis-regulatory elements (CREs), which are sequences that regulate the expression levels of target genes, sometimes across large genomic distances^5–7^. CREs include enhancers, promoters, and silencers and play key roles in establishing tissue-specific expression patterns. Genome-wide studies have extensively identified CREs and mapped them to their target genes^8–11^. Transcription factors (TFs) bind to these CREs, coordinating temporal and spatial expression profiles^12–16^. The human genome encodes approximately 1,600 TFs, with known target genes for many^17–19^. Regulation also occurs post-transcriptionally, involving factors such as microRNAs (miRNAs), that modulate mRNA stability and translation^20,21^. Over 2,000 human miRNAs have been annotated, and genome-wide interaction maps are available^22–25^. Gene regulation is also encoded within gene structure itself, as the length of untranslated regions (UTRs), coding sequences (CDS), and introns influences regulatory potential. Longer segments can accommodate more *cis*-acting elements, facilitating complex transcriptional and post-transcriptional control^26–29^. For instance, during early development, maternally supplied mRNAs retain long 3′ UTRs that support extensive post-transcriptional control, whereas zygotic mRNAs possess extended promoters to support the complex transcriptional programs required for initiating cell fate decisions^30^.

This diversity in expression patterns and regulatory mechanisms raises a fundamental question: how does regulatory architecture scale with expression pattern? Several models are possible. Broadly expressed genes may require complex regulatory mechanisms to maintain expression across the body. Conversely, tissue-specific genes might require tight regulation to ensure repression outside their intended context. Genes with intermediate tissue specificity might require intermediate levels of regulation or alternatively, the most elaborate control to balance selective activation and repression. Or, regulatory information content might be unrelated to expression patterns altogether and instead shaped primarily by other evolutionary or functional constraints.

To address this question, we conducted a genome-wide analysis of how gene expression patterns relate to multiple layers of regulatory control. Classic tissue specificity measures summarize where a gene is expressed but do not typically consider how expression patterns relate to the biological relationships between tissues. However, tissue relationships are meaningful: a gene expressed in three closely related immune tissues likely requires less regulatory information than one expressed in three unrelated tissues. To incorporate this dimension, we developed an ontogeny-aware, parsimony-based measure that estimates the minimal number of regulatory transitions required to explain a gene’s expression pattern across the tissue hierarchy. This measure ultimately quantifies the expression complexity, or regulatory demand, of a gene.

We applied this framework to examine the relationship between multiple regulatory features and the measured expression complexity of genes. We further tested whether this relationship behaves differently in selectively expressed versus ubiquitous genes, assessing whether the regulatory features act as binary ‘switches’ or quantitative ‘knobs’. We extended this analysis to explore how these patterns are shaped by evolutionary gene age. Collectively, this approach offers a quantitative framework for understanding how gene expression patterns are encoded by regulatory architecture and shaped evolutionarily.

## Results

### Tissue specificity spans a broad distribution with a substantial intermediate class

To quantify tissue specificity, we used the tau index^4^, a widely adopted measure that integrates both the number of tissues in which a gene is expressed and the variability of its expression levels across tissues. The tau values range from 0 (ubiquitous even expression across all tissues) to 1 (strict tissue specificity). A comparative evaluation of nine tissue specificity metrics identified tau as the most robust^31^, and it is widely used, e.g. in the Human Protein Atlas (HPA)^32,33^. We calculated tau values for 18,234 protein-coding genes from HPA expression data, derived from both bulk RNA-seq covering 40 tissues and single-cell RNA-seq data aggregated by 81 cell types^34^. Consistent with previous findings by HPA^34^, tau values computed from bulk data closely mirrored those derived from the single-cell data (Spearman’s correlation = 0.88, Supplementary Fig. 1A), indicating that tissue specificity is consistent across different biological resolutions.

The distribution of tau values exhibited a broad dynamic range, with peaks at both the low and high ends (Fig. 1A). While most genes exhibited either broad expression (low tau) or strict tissue specificity (high tau), a substantial subset fell in the intermediate range. By coloring the bar plot according to the number of tissues or cell types a gene is expressed in, we found that tau is strongly, but not exclusively, determined by expression breadth. This is further illustrated by plotting tau against the number of tissues (Supplementary Fig. 1B), revealing that broadly expressed genes can still span a wide range of tau values. Notably, genes with tau between 0 and ∼0.5 are expressed in essentially all tissues, indicating that roughly half of the tau range reflects expression level variation rather than number of tissues. Values near zero indicate uniform expression across tissues, whereas values approaching 0.5 reflect increasing unevenness; tau values above ∼0.5 correspond to progressively restricted expression, ultimately limited to one or a few tissues.

**Figure 1.**
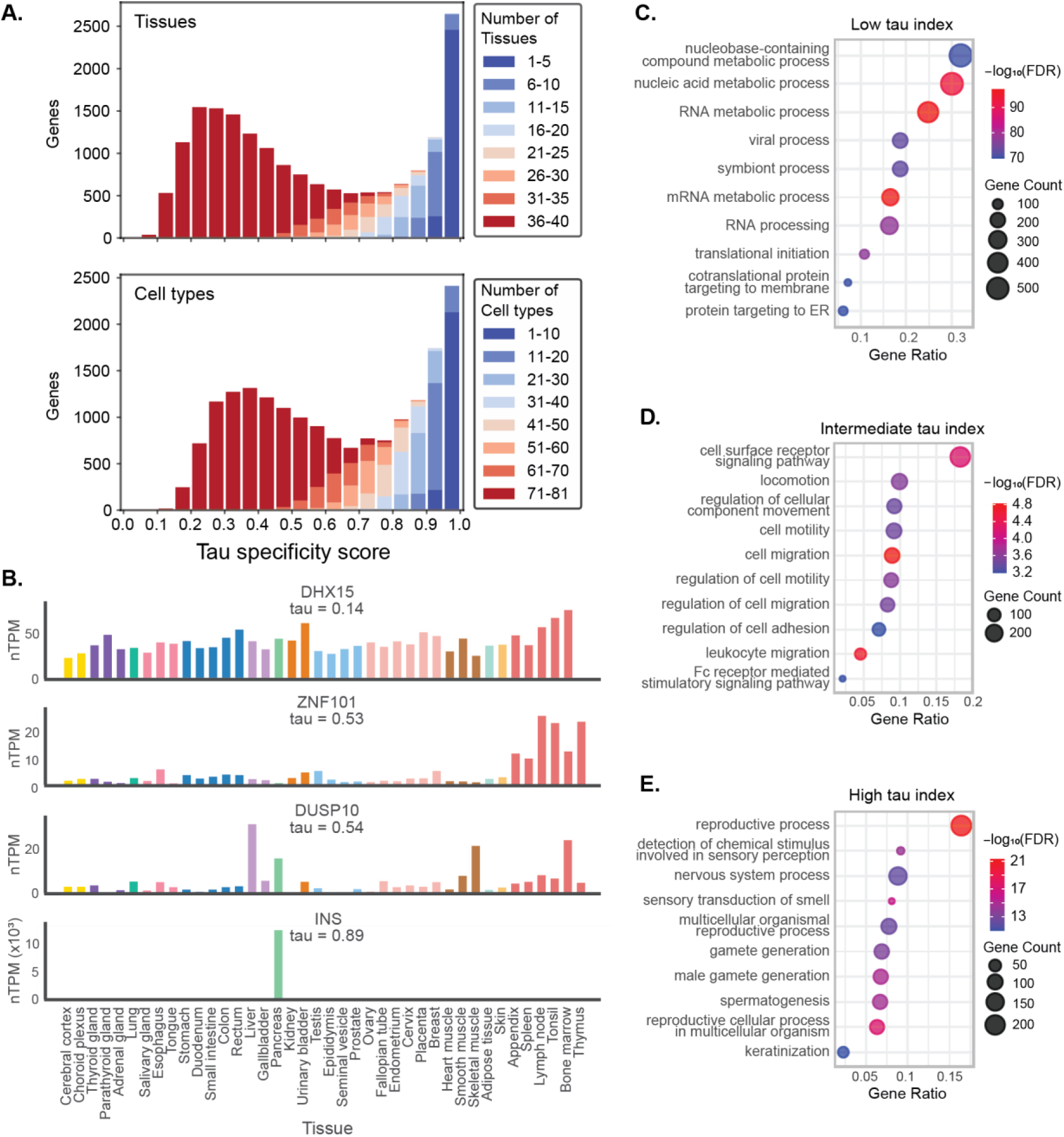
Tissue specificity of human protein-coding genes and associated functional enrichments. **(A)** Distribution of tau scores, calculated from bulk RNA-seq tissue samples (top) and single-cell RNA-seq data aggregated by cell type (bottom), for 18,325 human protein-coding genes. Tau ranges from 0 (ubiquitous, even expression) to 1 (strict tissue specificity). Bars are colored by the number of tissues or cell types in which each gene is expressed, illustrating the relationship between tau values and expression breadth. **(B)** Bar plots showing tissue expression profiles of representative genes spanning the tau spectrum. Tissues are color-coded by groups with shared functional characteristics, as defined by the HPA. Y-axis values indicate normalized gene expression (nTPM) across 40 tissues. **(C-E)** Bubble plots showing ranked GO term enrichment based on tau values: **(C)** Genes broadly expressed (low tau). **(D)** Genes with intermediate tissue specificity (tau = 0.5) **(E)** Highly tissue-specific genes (high tau). The x-axis represents the gene ratio, bubble size indicates gene count, and bubble color shows the –log₁₀ adjusted p-value.

To illustrate the range of expression profiles captured by the tau index, we display representative genes spanning the spectrum of tissue specificity. *DHX15*, an RNA helicase, exhibits uniform expression across all tissues and a low tau value (0.14), consistent with its housekeeping function. At the opposite extreme, the gene encoding insulin (*INS*) is predominantly expressed in the pancreas and shows a high tau value (0.89), exemplifying tissue specificity. Intermediate specificity is more heterogeneous, as demonstrated by two genes with tau values ∼0.5 (Fig. 1B). *ZNF101,* a zinc-finger protein involved in transcription regulation, is expressed at low levels across most tissues but shows elevated expression in immune-associated tissues such as lymph nodes, tonsils, and thymus. In contrast, *DUSP10*, which encodes a phosphatase that regulates MAP kinase activity, is also broadly expressed at low levels, but exhibits relatively high expression in a functionally diverse set of tissues including liver, pancreas, skeletal muscle, and bone marrow. While both genes share similar tau values, their expression patterns differ in the degree of tissue-related coherence. These examples highlight the diversity of expression patterns across the tau spectrum, with intermediate values capturing both tissue-coherent and functionally dispersed expression profiles.

To characterize biological processes associated with these gene-expression patterns, we performed ranked enrichment analyses using GOrilla^35^, ranking genes by tau values. Genes with low tau values were enriched for fundamental cellular functions, such as RNA metabolism and translation initiation, consistent with their housekeeping roles (Fig. 1C). Genes with high tau values showed significant enrichment for specialized processes, including sensory detection (olfaction), reproduction (spermatogenesis), and keratinization, matching tissue-specific functions (Fig. 1E). Genes with intermediate tau values were enriched for processes such as cell-migration and immune-related processes (Fig. 1D); albeit with lower statistical significance. This lower significance suggests that intermediate-specificity genes represent a biological phenomenon, yet they comprise a more functionally heterogeneous group.

These results collectively highlight a wide continuum of tissue specificity, motivating the question of how regulatory complexity scales across this spectrum.

### CRE abundance peaks at intermediate levels of tissue specificity

How might regulatory architecture relate to tissue specificity? To reveal this relationship, we first examined the distribution of regulatory elements across the tissue specificity spectrum. We began with CREs because they constitute the primary regulatory elements controlling spatiotemporal and cell-type-specific gene expression in metazoan genomes and are extensively annotated across diverse tissues. The number of CREs linked to a gene can serve as a genome-wide, quantitative proxy for the diversity of *cis*-regulatory inputs needed to implement its expression pattern.

Candidate CRE (cCRE) annotations and predicted gene-cCRE associations were obtained from the GeneHancer database^10^, which integrates data from multiple sources and assigns confidence scores to cCREs and their target links. We used both the full dataset (419,020 cCREs) and the more stringent “Double Elite” subset (122,815 cCREs), which includes only high-confidence cCRE-gene pairs. The main figures show results based on the full set, while Supplementary Fig. 2 presents the Double Elite analyses, with results consistent across both thresholds (see Methods).

We grouped genes into tissue specificity (tau) bins and in each bin, calculated the mean number of linked cCREs per gene. We observed an inverted U-shaped relationship. Genes with intermediate tissue specificity exhibited the highest mean cCRE counts, while both broadly expressed and highly tissue-specific genes have fewer cCREs (Fig. 2A). This pattern indicates that genes with intermediate tissue specificity require the most diverse cCRE inputs to achieve selective and context-dependent expression. Motivated by this finding, we sought a theoretical framework that could account for this non-monotonic relationship.

**Figure 2.**
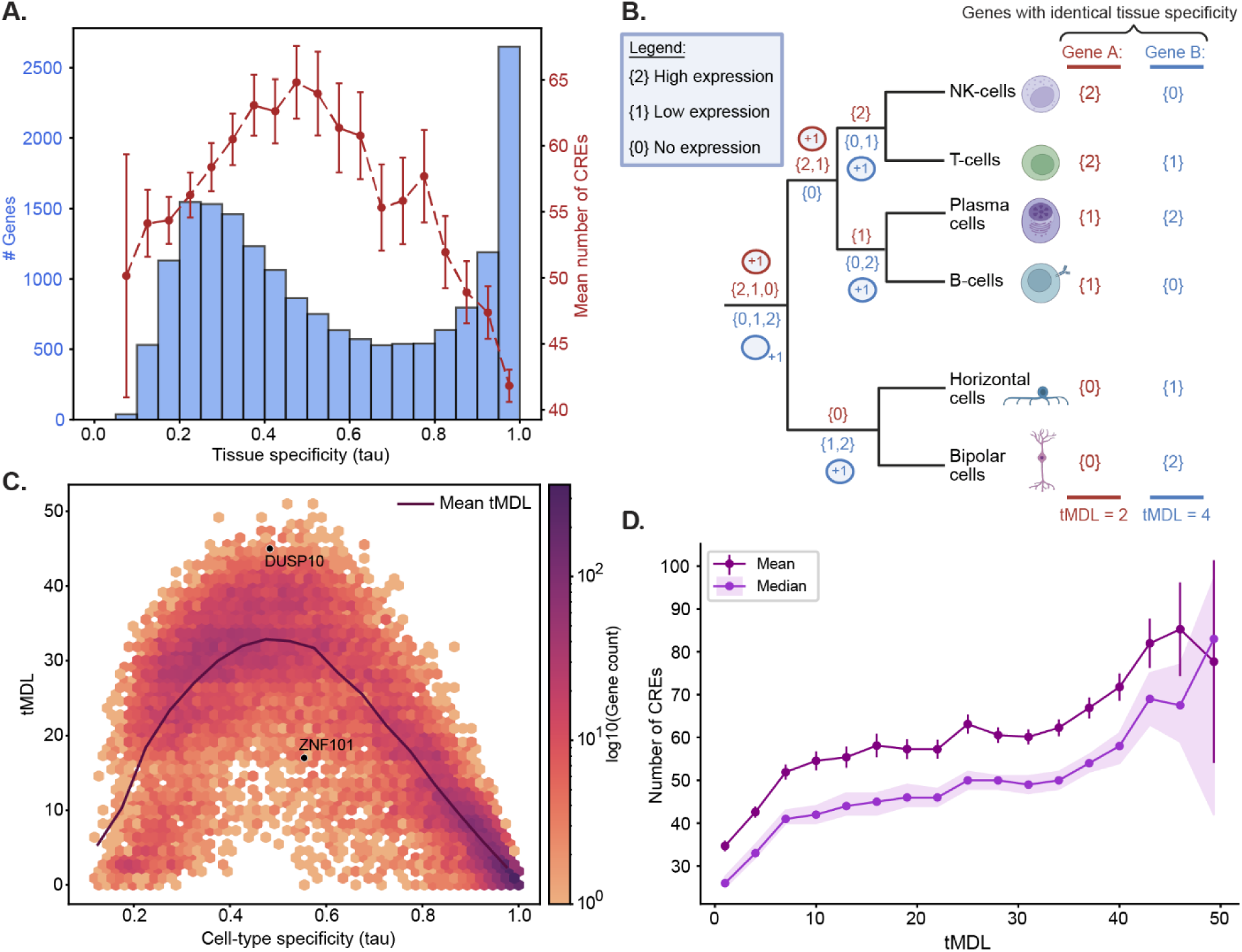
tMDL captures regulatory information content beyond tissue specificity. **(A)** Histogram of tau scores calculated from bulk RNA-seq data (blue bars, left y-axis), with overlaid line plot (red dashed line, right y-axis) indicating the mean number of CREs linked to genes within each tau bin, with error bars representing the 90% confidence interval (CI) of the mean. **(B)** Schematic of a simplified example illustrating how the tMDL is calculated using Fitch’s parsimony algorithm. Two hypothetical genes (Gene A, red; Gene B, blue) have identical tau values but differ in the distribution of their expression across a hierarchical tree of cell types. Leaf nodes show discretized expression levels: {0} for no expression, {1} for low expression, and {2} for high expression. Internal nodes are labeled with the possible expression states consistent with the expression patterns of their descendant cell types. Regulatory transitions are indicated along branches as “+1,” with each +1 representing one required expression-level change, and the total number accumulates along the path. The total number of transitions across the tree defines the tMDL score, shown at the bottom for each gene. Made with Biorender. **(C)** Hexbin plot showing the relationship between tau index and tMDL across all human protein-coding genes. Here, tau represents cell-type specificity. Both tau and tMDL are calculated with single-cell RNA-seq data aggregated by cell type. Each hexagon represents the genes within that region of the plot, with color indicating local density. Two example genes from Figure 1B. are marked with black dots, to demonstrate genes with similar tissue specificity but different tMDL. **(D)** Line plot showing the mean (purple) and median (violet) number of CREs per gene, computed in bins along ranked tMDL values using a sliding window approach (i.e. genes were ordered by tMDL, and the mean and median were calculated within fixed-size windows that slide across the ranked list). Error bars and shaded areas represent 90% CIs for mean and median, respectively.

### Minimum Description Length (MDL) as a framework for understanding gene regulatory architecture

The observation that cCRE abundance peaks in genes with intermediate tissue specificity suggests a non-monotonic relationship between tissue specificity and regulatory information content per gene. To explain this pattern, we turned to a principle from information theory called the Minimum Description Length (MDL). MDL states that the best representation of a system is the one that minimizes the amount of information required to describe it^36,37^. In essence, this frames gene regulation as a problem of information compression: how to encode a complex expression pattern concisely. Applied to gene regulation, this principle predicts that genes with simple expression patterns should require minimal regulatory architecture, while those with more complex patterns should require more elaborate control mechanisms.

For example, housekeeping genes may follow the simple rule “express in all tissues,” and tissue-specific genes follow “express only in tissue X.” Both represent concise regulatory instructions. In contrast, a gene that must be expressed in a subset of tissues, for instance “express in tissues A, D, and F but not or lowly in B, C, or E”, generally requires a more complex regulatory program. According to the MDL principle, regulatory information content should therefore follow an inverted U-shaped relationship with tissue specificity. Genes at either extreme with a short MDL describing their expression pattern should require simpler regulatory mechanisms, with lower information content in their regulatory program than those with intermediate tissue specificity.

### A maximum parsimony-based measure estimates MDL

While the tau index effectively captures expression distribution across tissues, it is only a proxy for estimating the regulatory information content predicted by MDL. The tau index does not account for the hierarchical or ontogenetic relationships between tissues. For example, both *ZNF101* and *DUSP10* have intermediate tau values (Fig. 1B), yet their expression profiles differ markedly in their tissue coherence. *ZNF101* is highly expressed in functionally related immune tissues such as lymph nodes, tonsils, and thymus, suggesting the potential for a shared regulatory program. In contrast, *DUSP10* shows elevated expression in a more dispersed set of tissues which are not related in expression programs. Although tau treats these genes as equivalent in specificity, their regulatory demands likely differ, with *ZNF101* requiring fewer developmental switches and thus a shorter MDL, while *DUSP10* may necessitate multiple independent regulatory instructions. The relationships between all tissues or cell types in the body can be depicted by a hierarchical tree of relatedness, approximated by similarity in expression programs^38^ (though exceptions can arise from convergent expression among developmentally distinct tissues). This underlying organization of tissues underscores the need for a tree-aware measure of regulatory demand that captures expression distribution in the context of this hierarchical structure.

Another computational biology domain that captures similarities based on hierarchical trees is phylogenetics. In evolutionary trees, maximum parsimony algorithms are used to infer the traits of ancestral species and to estimate the number of changes (e.g., nucleotide substitutions) along each branch. We recognized that the phylogenetic and ontogenetic problems are related, as both involve inferring the number of events (mutations in evolution or gene expression changes during development, respectively) that occur along each branch of a tree from a common node. This conceptual connection between phylogenetic methods and ontogeny has been explored in previous work that applied phylogenetic tools to expression-based cell type trees to gain developmental and functional insights^39–42^. Here we accordingly adapted a maximum parsimony approach from phylogenetics to quantify a gene’s regulatory demands. By analogy to phylogenetics, we constructed a hierarchical tree of tissue/cell type relationships using the genome-wide gene expression profiles and hierarchical clustering (see Methods). We then adapted Fitch’s algorithm^43^, originally developed to infer the minimum number of mutations required to explain sequence variation across a phylogenetic tree. In our framework, this algorithm is used to assess the minimal number of expression-level transitions needed to produce a given expression pattern, where a transition is defined as a change in discretized expression bin. The resulting score, referred to as the tree-aware MDL (tMDL), reflects the estimated minimal regulatory load required for a gene’s expression program while accounting for the relatedness of cell types, (see Fig. 2B, and Supplementary Fig. 3 for method’s robustness analyses). We used cell type–level data to build the tree, as single-cell-derived profiles provide a cleaner estimate of biological relationships than bulk tissue data, which can be confounded by compositional bias.

Higher tMDL scores indicate that more independent regulatory changes are required to produce the observed expression profile. Genes with high tMDL scores are generally expressed in several tissues that are not clustered together on the hierarchical tree. For instance, *ZNF101* and *DUSP10*, that both show intermediate tau values, differ markedly in their tMDL. *ZNF101* has a tMDL of 17, among the lowest observed at this tau level (Fig. 2C), because its high expression is restricted to closely related immune cell types. In contrast, *DUSP10* has a tMDL of 45, one of the highest for genes with similar tau, due to its high expression across a set of distantly related cell types. This illustrates how tMDL distinguishes between expression profiles that appear similar in breadth and tau but differ in the complexity of tissue-specific regulation.

Figure 2C shows, as predicted by the MDL principle, that genes with intermediate tissue specificity (tau ≈ 0.5) showed the highest tMDL, indicating they require the most regulatory information content. Notably though, genes in this intermediate range also exhibited considerable variation in their tMDLs, suggesting there are diverse levels of regulatory demands among genes with similar expression breadth and tissue specificity. This highlights how combining tau with tMDL provides a richer, more comprehensive view of regulatory architecture.

Genes with high tau (tissue-specific) showed low tMDL, near one, reflecting the simple logic of being “off” in most tissues and “on” in only one or a few, often requiring just a single regulatory transition. Genes with low tau (broadly expressed) generally had higher tMDLs than tissue-specific genes, but lower than many intermediate-tau genes. As tau approaches zero, tMDLs tend to decrease, consistent with the expectation that uniform expression requires fewer regulatory transitions. Nonetheless, broadly expressed genes also showed tMDL variability, reflecting differences in how consistent expression levels are maintained across tissues.

### Diverse regulatory features align with MDL framework

We next tested whether specific layers of gene regulation scale with complexity as predicted by MDL. We examined four regulatory features: cCREs, TFs, miRNAs, and lengths of various gene structural elements, assessing their relationship with regulatory demand (tMDL) and tissue specificity (tau).

### cCREs

As seen in Figure 2A, cCRE count per gene peaks among genes with intermediate tissue specificity. We asked whether cCRE count also scales with tMDL, thereby reinforcing its value as a proxy for regulatory information content. To test this, we performed a sliding window analysis of cCRE counts per gene, calculating the mean and median cCRE count across fixed intervals of increasing tMDL. This analysis revealed a clear trend: genes with higher tMDL scores tended to have more cCREs (Fig. 2D). We conclude that genes with more transitions in expression state within the tree require more inputs from cCREs.

### Transcription factors

Given the central role of TFs in modulating gene expression, we next examined how TF targeting relates to tissue specificity and regulatory information content. TF-target gene relationships were obtained from the hTFtarget database, which compiles experimentally supported TF binding events from ChIP-seq datasets and motif predictions, assigning targets when binding occurs in promoter regions^44^.

While cCRE count peaked among genes with intermediate tissue specificity, TF targeting followed a distinct pattern. Broadly expressed genes were targeted by the most TFs (Figure 3A). Interestingly, this increase in TF count plateaued around tau ≈ 0.4, suggesting that once a gene is widely expressed across all tissues, additional TFs provide diminishing returns. This trend differs from the cCRE pattern, suggesting that broadly expressed genes may require a more extensive *trans*-regulatory network to ensure robust and high expression across diverse cellular environments^45^. Genes with restricted expression breadth rely on more limited TF input, requiring activation in only a narrow set of conditions or cell types. We note that TFs can act as activators, repressors, or possess dual roles depending on cellular context and target gene^46–48^.

**Figure 3.**
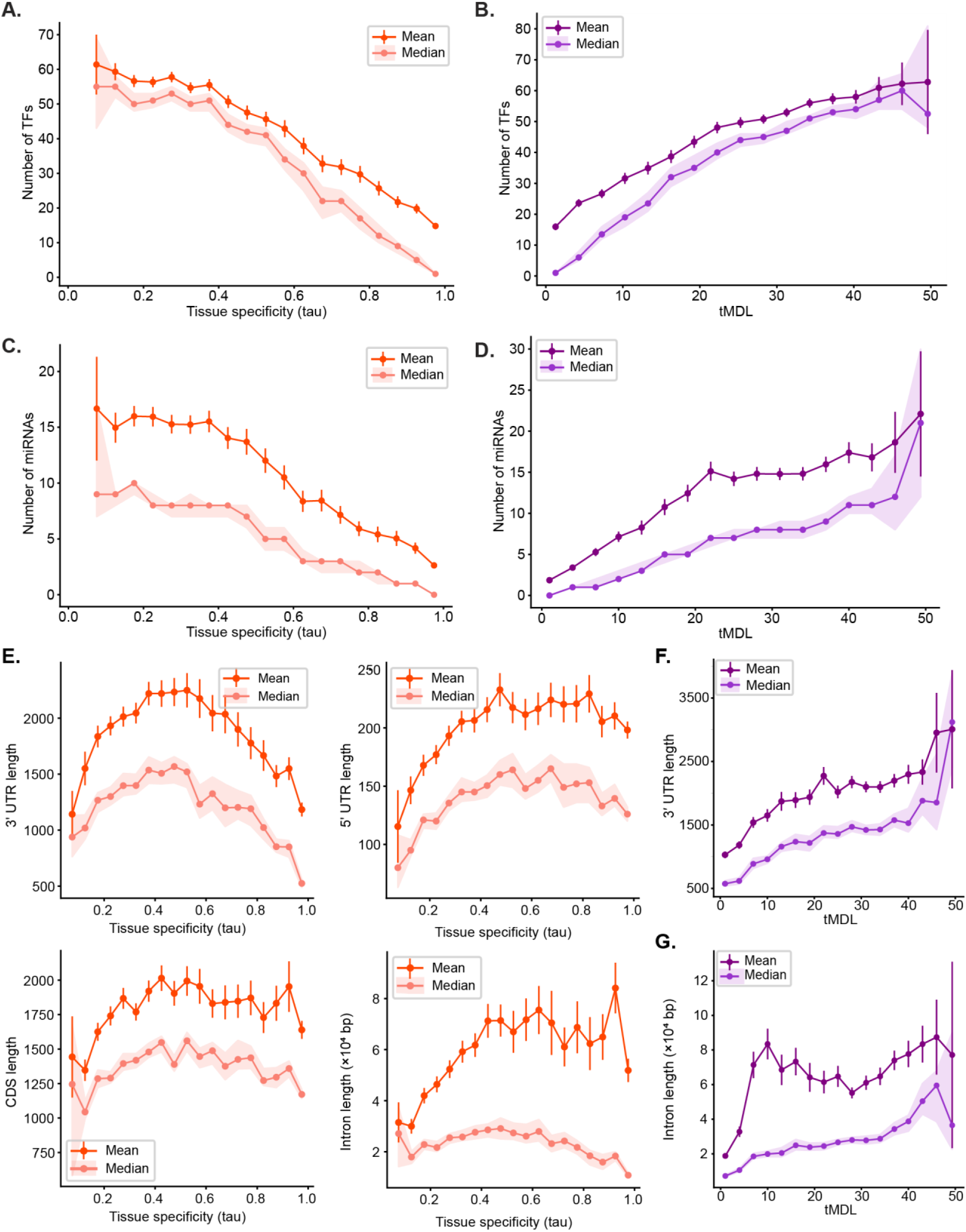
Transcriptional and structural regulatory features scale with regulatory demand (tMDL). **(A–B)** Mean and median TF counts per gene were computed across sliding windows of ranked tau (A) or tMDL (B) values. **(C–D)** Mean and median miRNA counts per gene across bins of tau (C) and tMDL (D). **(E)** Gene structure features across tau: 3′ UTR, 5′ UTR, CDS, and intron lengths. **(F–G)** 3′ UTR length (F) and intron length (G) across bins of tMDL. In all panels, dark lines with error bars (90% CI) represent mean values and light lines with shaded area (90% CI) represent median values. Orange hues correspond to tau-based plots, and purple hues to tMDL-based plots. Tau is analyzed using equal-width bins across its value range; tMDL using fixed-size sliding windows over ranked genes.

Despite these differences regarding tissue specificity, we observed a strong positive trend between TF number and tMDL (Figure 3B). Genes with more complex expression patterns, as captured by higher tMDLs, tended to be targeted by more TFs. Although TF number declines steadily with increasing tissue specificity, its robust progressive increase with tMDL suggests that TF targeting still reflects overall regulatory burden, capturing aspects of regulatory complexity that go beyond tissue specificity alone. These findings show that, like CRE count, TF targeting scales with regulatory demand as captured by tMDL.

### microRNAs

We next turned to post-transcriptional regulation, focusing on miRNAs. Experimentally detected miRNA-target interactions were obtained from TarBase, a comprehensive, manually curated database that catalogs over 2 million unique miRNA-gene pairs supported by diverse experimental methods^49^. Similar to TF targeting, we found that broadly expressed genes harbored the highest number of targeting miRNAs, while tissue-specific genes contained the fewest (Figure 3C). When compared with tMDL, the number of targeting miRNAs showed a strong positive slope (Figure 3D), meaning genes with more complex expression patterns tended to be targeted by a greater number of miRNAs. This trend demonstrates that genes under higher regulatory demand also engage more extensively with post-transcriptional mechanisms, aligning with the MDL framework’s prediction. It closely parallels the trend for TFs, which have been shown to co-regulate shared targets with miRNAs and even regulate each other in integrated network motifs^50^. However, TF and miRNA counts showed a low correlation (Supplementary Fig.4), indicating their similar scaling with tMDL is not due to a dependency between them. The progressive increase in both miRNA and TF targeting with tMDL suggests that both transcriptional and post-transcriptional trans-regulatory inputs scale with overall regulatory demand, even when not directly reflected by tissue specificity.

### Gene structure

We next examined how gene structural features (i.e. CDS, intron, and UTR lengths) relate to tissue specificity and tMDL. Prior studies have linked gene structure to regulatory potential and mRNA stability^27,30,51–54^. Additionally, broadly expressed housekeeping genes are known to be typically shorter; an observation thought to reflect selection against the energetic cost of long transcripts^55^. Under the MDL framework, tissue-specific genes are also expected to be relatively short, since regulatory information may be embedded within the gene’s structure too. In contrast, genes with more complex expression patterns may encode additional regulatory instructions within their structure, resulting in increased length.

Consistent with this, genes with intermediate tissue specificity showed the longest CDS, introns, and UTRs (Fig. 3E). While some features displayed a more asymmetrical inverted U-shape, particularly with broadly expressed genes often being the shortest, the 3′ UTR length showed the most pronounced trend of peaking among genes with intermediate specificity.

We next asked whether these structural features correlate with tMDL. Both 3′ UTR and intron lengths increased with tMDL, indicating that genes requiring more regulatory state transitions tend to have longer non-coding regions (Fig. 3F-G). This aligns with their known regulatory roles: 3′ UTRs often serves as hubs for post-transcriptional control, including microRNA, RBP interactions, and other regulatory elements, while introns can harbor CREs, miRNA binding sites, and additional regulatory elements^56–58^. Among all features, 3′ UTR length showed the strongest relationship with both tau and tMDL, highlighting its regulatory importance. Although longer 3′ UTRs offer more space for miRNA binding, the correlation between miRNA count and 3′ UTR length was only moderate (Supplementary Fig. 4), indicating that UTR length captures additional regulatory complexity. In contrast, CDS and 5′ UTR lengths showed little to no correlation with tMDL, consistent with the 5′ UTR’s primary role in translation initiation rather than expression regulation^59^, and with prior observations that 5′ UTR length does not contribute to tissue specificity^60^.

Together, these findings highlight gene structure, particularly 3′ UTRs, as a proxy for embedded regulatory information content. Longer genes appear to accommodate more intricate regulatory mechanisms, consistent with MDL predictions.

### Quantitative framework distinguishes knob-like and switch-like regulatory mechanisms across gene expression regimes

Genes with tau values below ∼0.5 are broadly expressed across all tissues and thus show variation in their relative expression levels rather than presence or absence in each tissue. These genes are expected to require fine-tuned, quantitative control, hence refer to here as “knob-like” regulation. In contrast, genes with higher tau values are expressed in a more restricted set of tissues and thus require more selective activation or repression, suggestive of “switch-like” regulation. To explore how different regulatory mechanisms contribute to these distinct regimes, we divided genes into two groups (Fig. 4A): those expressed in ≥90% of cell types (ubiquitous) and those expressed in fewer than 90% (cell-type-selective). We then asked how regulatory features and tMDL scaled within each group, to assess whether different regulatory strategies support distinct expression regimes.

**Figure 4.**
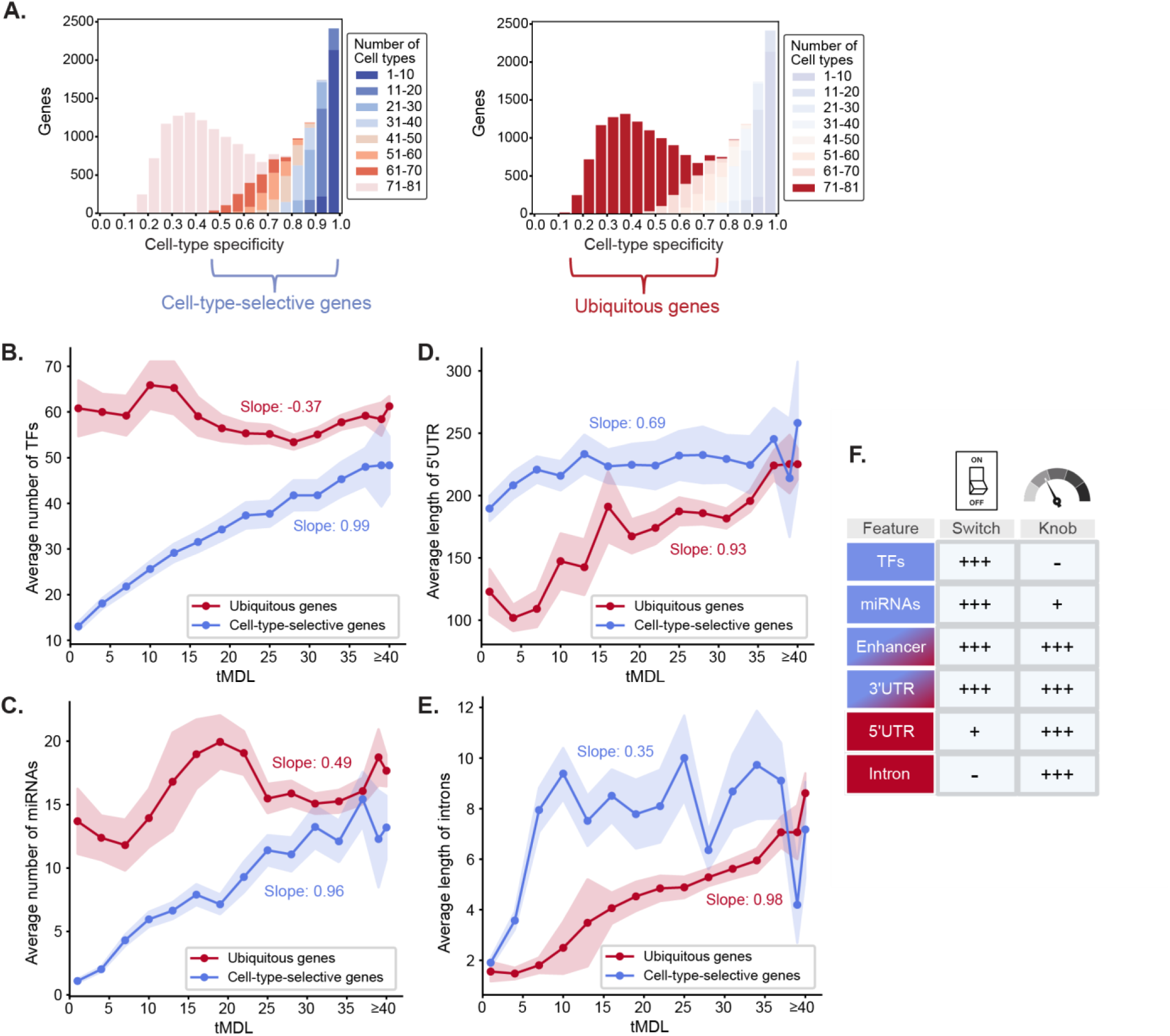
Distinct regulatory features scale with tMDL in cell-type selective versus ubiquitous genes. **(A)** Histograms showing the distribution of tau values, highlighting the two expression regimes: cell-type selective genes (expressed in <90% of tissues; left) and ubiquitous genes (≥90% of tissues; right). Colored bars indicate the number of tissues in which each gene is expressed; transparent bars denote genes outside the respective group. **(B–E)** Mean feature values were calculated across sliding windows of ranked tMDL, separately for ubiquitous (red) and cell-type selective (blue) genes. Slopes were obtained from linear models fit to windowed means after standardizing both variables (zero mean, unit variance) to allow comparison across features. Shaded areas represent 90% CI. To avoid sparse windows, all genes with tMDL ≥ 40 were grouped into a final bin. **(B)** Number of TFs per gene. **(C)** Number of miRNAs per gene. **(D)** 5′ UTR length. **(E)** Intron length. **(F)** Table summarizing regulatory features according to their association with tMDL in tissue-selective versus broadly expressed genes, representing switch-like and knob-like regulatory regimes, respectively. Symbols indicate the strength of the observed trend (based on linear regression slope): ‘+++’ = strong, ‘++’ = moderate, ‘+’ = weak, ‘–’ = none or negligible.

CRE abundance showed similarly robust correlations with tMDL in both broadly and selectively expressed genes (Supplementary Fig. 5), highlighting their universal importance across regulatory strategies, both knob-like and switch-like. However, TF targeting demonstrated strikingly different dynamics in the knob and switch regulated genes (Figure 4B). In tissue-selective genes, TF numbers increased greatly with rising tMDL, consistent with their important roles in discrete activation or repression events across specific tissues. Conversely, broadly expressed genes displayed high and relatively stable TF targeting, with a slight decline across tMDL. This finding suggests that broadly expressed genes utilize a largely constant set of TFs, indicating that additional fine-tuning regulatory demands in these genes likely arise through alternative mechanisms rather than changes in TF number, potentially involving differential TF activity, co-factor interactions, CREs, or other regulatory inputs.

miRNA regulation patterns paralleled those observed for TFs (Figure 4C). Tissue-selective genes exhibited a strong positive relationship between miRNA count and tMDL, reinforcing miRNAs’ critical role as regulatory switches. Broadly expressed genes showed moderately increased miRNA targeting with rising tMDL, though this relationship was less steep. Thus, while miRNAs are broadly employed, they scale particularly strongly with regulatory complexity in tissue-selective genes, highlighting their dominant role as switch-like regulators.

Gene structural features revealed additional complexity. Lengths of 3′ UTR regions increased consistently with tMDL in both expression regimes, highlighting their universal regulatory role (Supplementary Fig. 5). However, introns and 5′ UTR lengths exhibited regime-specific behavior: both increased significantly with tMDL in broadly expressed genes, but showed weaker trends in cell-type selective genes (Figure 4D,E), indicating a specialized role in fine-tuning regulatory complexity. This suggests that introns and 5′ UTRs contribute to fine-tuning in broadly expressed genes, but are not a major component of switch-like regulation in tissue-selective genes. As gene length is known to be constrained by energetic cost, particularly in highly expressed genes, the scaling of intron and 5′ UTR length with tMDL in broadly expressed genes may reflect selective pressure to maintain compactness except where additional regulatory content is required.

Together, these findings support a model in which distinct regulatory strategies underlie broadly versus selectively expressed genes. While some features, such as CREs and 3′ UTRs, scale with regulatory complexity across both regimes, others show regime-specific patterns. These distinctions are summarized in Figure 4F, which classifies each regulatory feature according to its predominant role in switch-like versus knob-like regulation based on our observed patterns.

### Chromosome X as a potential case of chromosome-level MDL compression

Thus far, our analyses have examined the regulatory demands of individual genes across the genome. However, we observed that these principles may also extend to higher-order genomic organization. In agreement with previous studies, we found that genes located on the X chromosome (chrX) exhibit a significant bias toward tissue-specific expression^61,62^. Specifically, 25.3% of chrX genes exhibited high tissue specificity (tau ≥ 0.95), compared to 14.5% genome-wide (Figure 5A).

**Figure 5.**
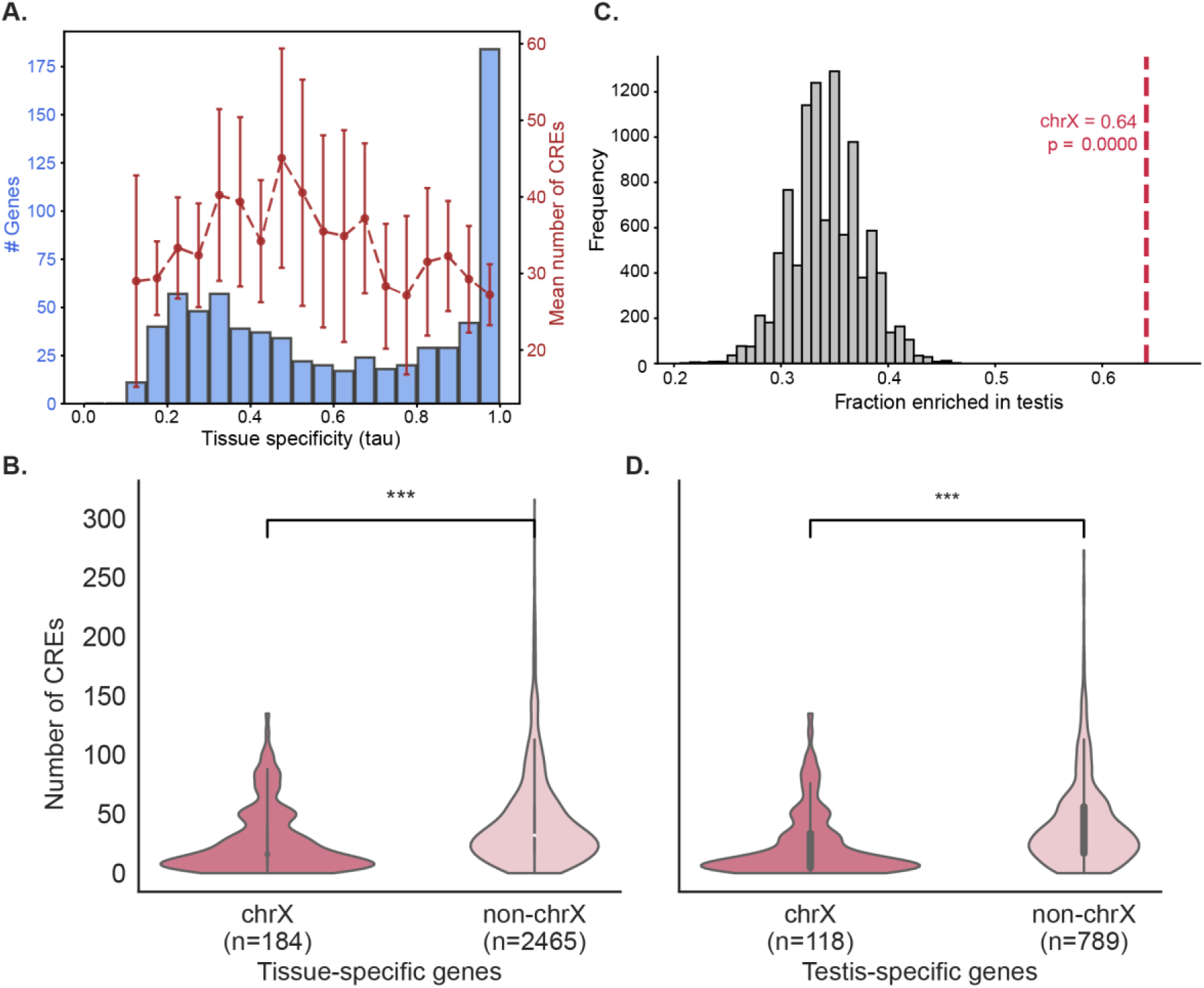
Tissue-specific genes on chrX have reduced CRE counts and are enriched for testis-specific expression. **(A)** Histogram showing the number of genes on chrX across bins of tissue specificity (tau). Overlaid in red is the mean number of CREs per gene in each tau bin. Error bars represent 90% CI of the mean. **(B)** Violin plot comparing the number of CREs per gene in tissue-specific genes on chrX (tau ≥ 0.95, dark pink) versus tissue-specific genes on the rest of the chromosomes (light pink), showing significantly lower CRE counts on chrX (Mann–Whitney U test, *p* = 5.2 × 10⁻¹¹). **(C)** Null distribution showing the fraction of testis-specific genes (defined by dominant tissue of highest nTPM and tau ≥ 0.95) across 10,000 randomly sampled gene sets matched for tissue specificity. The observed chrX proportion (0.64) is shown as a red dashed line, with its empirical p-value, demonstrating a significant enrichment of testis-specific genes on chrX beyond what is expected by chance. **(D)** Violin plot comparing the number of CREs for testis-specific genes on chrX (dark pink) versus testis-specific genes on the rest of the chromosomes (light pink), showing significantly lower CRE counts on chrX (Mann–Whitney U test, *p* = 4.4 × 10⁻^10^).

Focusing on these tissue-specific chrX genes, we found that they are associated with a significantly lower number of CREs compared to all tissue-specific genes across the genome (Figure 5B). This indicates that the regulatory demand for tissue-specific genes on chrX is lower than expected, even given their restricted expression patterns.

To understand this further, we asked whether these genes are enriched for specific tissues. As previously observed^63–65^, we found a strong enrichment for testis-specific expression: approximately 65% of the tissue-specific genes on chrX are testis-specific. To evaluate whether this enrichment exceeded expectations, we generated 10,000 random gene sets matched for tau distribution. These typically included only 30-40% testis-specific genes, confirming a significant overrepresentation of testis-specific genes on chrX (Figure 5C). Because the majority of chrX tissue-specific genes are testis-specific, we asked whether low CRE counts are a general feature of testis-specific expression. However, testis-specific genes on chrX had significantly fewer CREs than those on other chromosomes, suggesting a unique regulatory pattern on chrX (Figure 5D).

These findings reveal that chrX harbors a disproportionately high number of testis-specific genes with lower-than-expected CRE counts. This pattern supports that MDL principles may apply not only to individual gene expression patterns but also to higher-order genomic organization. From an MDL perspective, when many nearby genes share a simple expression rule, such as “express in testis”, the collective regulatory burden can be reduced, minimizing the description length required. Such compression at the chromosome level could reflect a broader organizational principle, potentially shaping how regulatory programs are spatially arranged across the genome. While this observation aligns with the MDL framework, alternative explanations for the low cCRE count on chrX are possible and warrant further investigation.

### Tissue specificity and regulatory information content vary across evolutionary age

Gene expression and regulation evolve over time, with newly emerged genes integrating into existing networks while older genes are maintained, or lost, across species. Comparative studies have shown that older genes often possess richer regulatory landscapes, including more alternative splice forms and TF binding sites than recently evolved genes^66–68^. However, because gene age is closely tied to tissue specificity (older genes are typically broadly expressed, younger genes more tissue-specific^69,70^), this relationship must be considered when evaluating age effects on regulatory architecture. Accordingly, we examined how evolutionary age relates to tissue specificity, regulatory features, and tMDL.

Gene ages were defined according to Thomas et al., who assigned each human gene to one of 19 phylostrata based on the most distantly related species in which an ortholog is detected, ranging from genes shared by all living organisms (phylostratum 1) to primate-specific genes (phylostratum 19) ^71^. Analysis of tissue specificity across age groups revealed a gradual transition, with each successive phylostratum showing progressively higher tissue specificity (Figure 6A). This pattern agrees with previous reports^69,70^.

**Figure 6.**
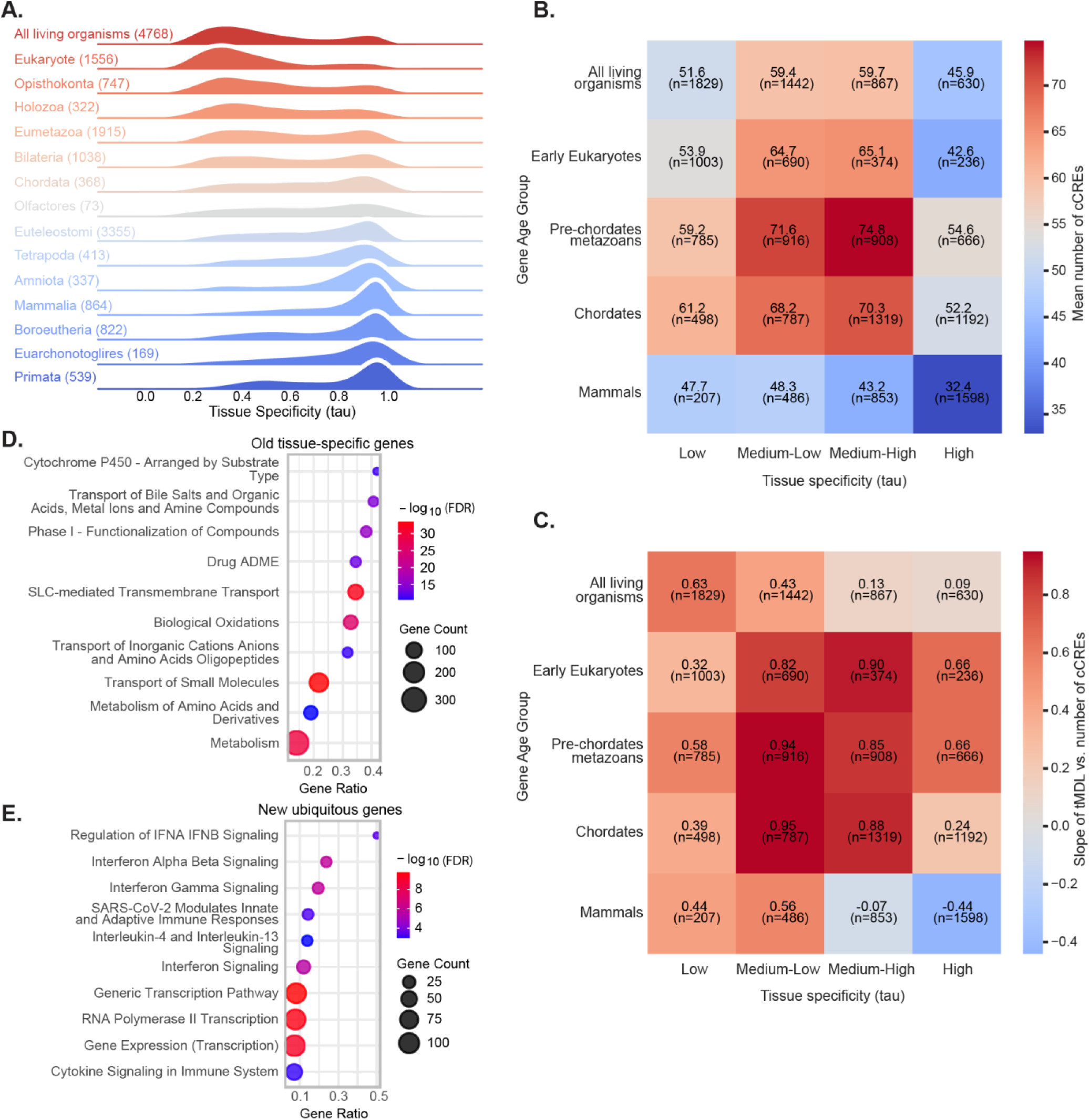
Tissue specificity and regulatory architecture vary across evolutionary gene age. **(A)** Smoothed density plots showing the distribution of tau values across evolutionary age groups. Genes are stratified by phylostrata, with the evolutionary age of each group indicated on the y-axis. Colors are used to distinguish age groups but carry no additional meaning. **(B)** Heatmaps displaying the mean number of cCREs for gene groups stratified by both evolutionary age (y-axis) and tissue specificity (x-axis, binned by tau values). **(C)** Heatmaps showing the normalized slope from sliding window analysis of cCRE number versus tMDL, with genes grouped by evolutionary age (y-axis) and tissue specificity (x-axis, binned by tau values). **(D–E)** Bubble plots showing the top 10 significantly enriched Reactome pathways for (D) genes with high tissue specificity (tau > 0.8) and ancient evolutionary origin (phylostrata 1–4) and (E) genes with low tissue specificity (tau < 0.4) and recent evolutionary origin (phylostrata 12–19). The x-axis represents the gene ratio, bubble size indicates gene count, and bubble color shows the –log₁₀ adjusted p-value.

Building on this, we next asked whether regulatory architecture also changes systematically with gene age, focusing on cCRE abundance across evolutionary groups. To control for the confounding effect of tissue specificity, we divided genes into four tau bins (low, medium-low, medium-high, high specificity) and five evolutionary age categories grouped from the original phylostrata classifications (“All living species,” “Early Eukaryotes,” “Pre-chordate metazoans,” “Chordates,” “Mammals”).

This analysis revealed that cCRE abundance consistently peaked in genes of intermediate evolutionary age across all tau levels, with these genes showing higher cCRE counts than both older and younger groups (Figure 6B). Thus, regulatory information content is shaped not only by tissue specificity but also across a gene’s evolutionary age. Within each age group, except for the most recent, cCRE count was highest among genes with medium-low or medium-high tissue specificity, consistent with our previous results.

We then asked whether the strength of the cCRE–tMDL relationship, meaning how strongly cCRE number scales with a gene’s regulatory demand, varies across evolutionary time. For each gene group, we calculated the slope of cCRE number versus tMDL. Genes of intermediate age exhibited the steepest positive slopes (Figure 6C). In contrast, the oldest (prokaryote-origin) and youngest (mammal-specific) genes showed flatter slopes, indicating a weaker relationship between cCRE usage and regulatory demand. Housekeeping genes with low tissue specificity consistently displayed shallow slopes across all age groups, possibly reflecting selective pressure to minimize regulatory burden in genes requiring stable expression.

While these findings establish a clear relationship between gene age, tissue specificity, and regulatory information content, some genes display interesting deviations from this pattern. Specifically, ancient genes with tissue-specific expression and young genes with already broad expression. To investigate whether these atypical expression patterns reflect underlying biological functions, we performed pathway enrichment analysis using Reactome annotations^72^. Ancient tissue-specific genes were enriched for pathways in xenobiotic metabolism, small molecule transport, and amino acid metabolism (Fig. 6D), suggesting roles in cell-autonomous detoxification, metabolic processing, and handling of endogenous and exogenous compounds. These genes may represent cell autonomous adaptation^73^. These functions, essential in unicellular organisms, have been conserved but are now localized to specialized organs in multicellular species. Tissue enrichment analysis revealed overrepresentation in testis, liver, and kidney (Supplementary Fig. 6), consistent with their metabolic and detoxification roles and the high transcriptional activity of the testis.

Recently evolved yet broadly expressed genes were enriched for transcriptional and innate immune pathways, including “Generic Transcription Pathway,” “RNA Polymerase II Transcription,” and interferon signaling cascades (Fig. 6E). Many of these genes belong to the rapidly evolving zinc finger (ZF) family^74,75^. Although sequence divergence classifies them as recently evolved, their broad expression and core functions suggest origins in ancient housekeeping genes. Rapid evolution may mask these deeper roots, while ancestral regulatory architecture enables their integration into essential gene networks.

Collectively, these results highlight that the relationship between gene expression complexity and regulatory architecture is modulated across evolutionary history, with the clearest MDL-consistent behavior emerging in genes of intermediate evolutionary age.

## Discussion

Our study presents a new conceptual framework for understanding regulatory information content in the human genome, grounded in the principle of MDL and maximum parsimony. By treating gene expression as an information compression problem where regulatory architecture encodes the rules for tissue-specific expression, we show that regulatory architecture follows predictable patterns of information content, shaped by both expression patterns and evolutionary history.

A key advance of this work is our maximum parsimony-based measure quantifying the minimal number of regulatory transitions needed for a gene’s expression profile across tissues or cell types. Unlike conventional tissue specificity measures such as tau, tMDL captures the hierarchical relationships between cell types and accounts for expression variation across both closely related and distantly related cell types. One important consideration is that tMDL depends on the choice of cell-type hierarchy. We used an expression-based tree because it captures similarities in transcriptional programs between cell types, and thus their underlying regulatory environments, making it well-suited for addressing questions of regulatory similarity. While cells from the same lineage are often transcriptionally similar^76^, such trees may not fully reflect developmental or differentiation relationships. Efforts to reconstruct complete developmental lineage trees of cells across the human body are advancing^77–79^ and once available, integrating them with expression-based hierarchies could further strengthen our framework.

While nearly all regulatory features scaled with our predicted regulatory demand (tMDL), only some features showed MDL-consistent trends when analyzed against tissue specificity (tau). *Cis*-regulatory features like cCRE number, intron length, and 3′ UTR length peaked among intermediate tau values, consistent with MDL and with prior reports^80–82^. In contrast, *trans*-acting features such as TFs and miRNAs increased monotonically with decreasing tau, plateauing among broadly expressed genes. This contrast reflects fundamental differences between *cis*- and *trans*-regulation: while *cis*-elements support modular, tissue-specific, and context-specific control, *trans*-regulators often act globally and are reused across many genes and tissues. The divergent patterns seen between tau and tMDL in *trans*-regulators highlight the limitations of using tissue specificity alone to infer regulatory information content. While tau effectively captures expression breadth and variability, it does not account for the hierarchical relationships between tissues or cell types, a key feature incorporated into the tMDL. This alignment with tMDL, even in the absence of a consistent trend with tau, highlights how tMDL more effectively captures regulatory demand encoded in gene expression patterns. Notably, pairwise correlations between these regulatory features were generally moderate to low (Supplementary Fig. 4), indicating that the observed trends are not simply due to dependencies among features.

Further, our analyses provide a quantitative framework for distinguishing “switch-like” versus “knob-like” regulatory strategies. Using our tMDL-based approach, regulatory features can be evaluated across expression regimes to determine their predominant mode. This framework could be extended to other regulatory layers or organisms for similar classification.

The relationship between expression patterns and regulatory information content is also shaped by evolutionary history. Genes of intermediate evolutionary age showed the highest regulatory information content, as reflected in their elevated cCRE content and alignment with MDL predictions, indicating that regulatory architecture follows a dynamic rather than linear trajectory. One possible explanation is that newly emerged genes, often tissue-specific, lack extensive cis-regulatory infrastructure and gradually acquire CREs as they integrate into regulatory networks. Over longer timescales, selective pressures on very ancient, often housekeeping genes may streamline regulation and reduce CRE abundance. In this view, intermediate-age genes may occupy a transitional phase: sufficiently evolved to have acquired diverse regulatory mechanisms but not yet constrained by the economizing pressures seen in the oldest genes.

While our results focus on patterns within a species, the regulatory landscape itself evolves as organisms acquire more cell types. An intriguing direction for future research is to examine how tissue specificity and regulatory information content shift across species with increasing cellular diversity, which could further support the MDL framework as a principle governing both intra-species regulation and macroevolutionary scaling. Equally compelling is the exploration of temporal dynamics, testing whether MDL principles apply to changes in regulatory information content during development within a tissue or organism, alongside the spatial patterns observed across tissues.

In conclusion, our results show that regulatory information content scales with the informational demands of a gene’s expression pattern, consistent with MDL predictions, and is further shaped by evolutionary history. While other factors such as environmental cues, developmental programs, and network context also contribute, expression complexity emerges as a major driver of regulatory burden. By integrating expression patterns, regulatory features, and evolutionary dynamics within an information-theoretic framework, we identify a fundamental organizing principle of gene regulation. Future work spanning cross-species comparisons, developmental time courses, and perturbation experiments could refine this model and clarify how regulatory complexity evolves. More broadly, our approach illustrates how information theory can provide a quantitative framework for understanding the forces shaping genome regulation over evolutionary timescales.

## Methods

### Expression datasets

Transcript-level RNA-seq data was downloaded from the HPA website (version 24.0) on December 25, 2024^33^. Bulk tissue data includes normalized transcripts per million (nTPM) values for 40 normal human tissues based on Ensembl v109 annotation. Single-cell derived data includes nTPM values across 81 annotated cell types from 31 tissues.

### Calculation of tissue specificity (tau) indices

We calculated the tau index separately for bulk and single-cell datasets to quantify the tissue specificity of each gene. The tau index ranges from 0 (ubiquitous expression) to 1 (highly specific expression) and was computed following the formula by Yanai et al.^4^:

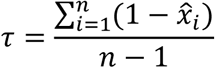

In this formula, *n* is the number of tissues or cell types, and *xᵢ* is the expression value in tissue or cell type *i*, normalized by the gene’s maximum expression across all *n* samples.

Expression values were first log-transformed (log(nTPM + 1)). After excluding genes that were not detected across any tissue or cell type, the tau index was computed for 18,234 protein-coding genes.

### Gene enrichment analyses

The Gene Ontology term enrichment was performed with the GOrilla web tool^35^. Separate ranked-list analyses were run for each tissue specificity class. For the high tau set, genes were ranked in descending order of tau; for the low tau set, in ascending order; for the intermediate tau set, by absolute deviation from tau = 0.5 (|tau – 0.5|, smallest to largest). Default parameters were used in single-ranked-list mode, and enrichment p-values were adjusted by the tool’s built-in Benjamini–Hochberg procedure.

The Reactome pathway enrichment was performed with the Enrichr web interface (Reactome 2024 library)^83^. Ancient tissue-specific gene sets were defined as those with tau ≥ 0.8 and assigned to the Holozoa or earlier evolutionary age groups (666 gene total). New ubiquitous genes were defined as those with tau ≤ 0.4 and assigned to the Euteleostomi or more recent age groups (1,316 genes total). P-values were corrected using the Benjamini–Hochberg method. Tissue-level enrichment for these same gene sets was assessed using the TissueEnrich web tool (https://tissueenrich.gdcb.iastate.edu/) with default settings^84^.

### cCRE annotations and gene-cCRE links

cCRE annotations and genec-CRE links were obtained from GeneHancer^10^, which integrates data from multiple sources and assigns confidence scores to predicted gene targets. We performed all analyses using both the full dataset (419,020 cCREs) and the high-confidence “Double Elite” subset (122,815 cCREs), in which both the enhancer itself and its link to the target gene are supported by at least two independent sources, providing stronger evidence for the regulatory association. In the full dataset, each gene was linked to an average of ∼55 enhancers, and each enhancer to ∼7 genes; in the Double Elite subset, each gene was linked to ∼7 enhancers, and each enhancer to ∼6 genes. Results were consistent across both sets; primary analyses used the full set, with Double Elite results shown in Supplementary Fig.2.

### Construction of cell-type similarity tree and calculation of tree-aware Minimum Description Length (tMDL) scores

Gene expression profiles (log-transformed nTPM, values < 1 set to zero) across 81 cell types were used to construct a cell-type similarity dendrogram. Pairwise correlations between cell types were computed using Spearman’s correlation coefficient, and a hierarchical clustering (Ward’s method) was performed on the correlation-based distance matrix (distance = 1 − Spearman correlation). The resulting linkage matrix was converted into a rooted dendrogram.

To quantify the tMDL for each gene, expression levels across cell types were discretized into six equally populated bins, with zero expression allocated to a separate bin. Using this discretization, Fitch’s parsimony algorithm^43^ was applied to the dendrogram, computing the minimal number of state transitions required to explain each gene’s binned expression pattern across cell types. The algorithm assigns possible expression states to internal nodes by working recursively from the leaves toward the root. At each internal node, the intersection of expression states from its descendant nodes is taken; if this intersection contains at least one common state, it is assigned to the node without incrementing the count. However, if there is no common state (i.e., the intersection is empty), the node is assigned the union of descendant states, and the count of required state transitions is incremented by one. The final sum of these increments across the dendrogram was reported as the tMDL score for each gene.

Unless otherwise specified, tau values were derived from bulk RNA-seq tissue samples from the HPA, to maintain consistency with prior large-scale studies and enable comparisons. In contrast, tMDL requires a hierarchy based on expression similarity, which at the tissue level can be confounded by shared or contaminating cell populations. For example, tissues with high blood or immune cell infiltration may appear transcriptionally similar due to expression from circulating cells rather than true similarity between the tissues. We therefore calculated tMDL using single-cell RNA-seq data aggregated by cell type, which mitigates these issues and produces a more biologically meaningful hierarchy. Gene expression patterns were highly concordant between bulk and single-cell datasets, and complementary analyses using tau from cell-type data and tMDL from tissue data produced overall similar trends (Supplementary Fig. 7), supporting the robustness of our conclusions.

### External data sources and their usage in this study

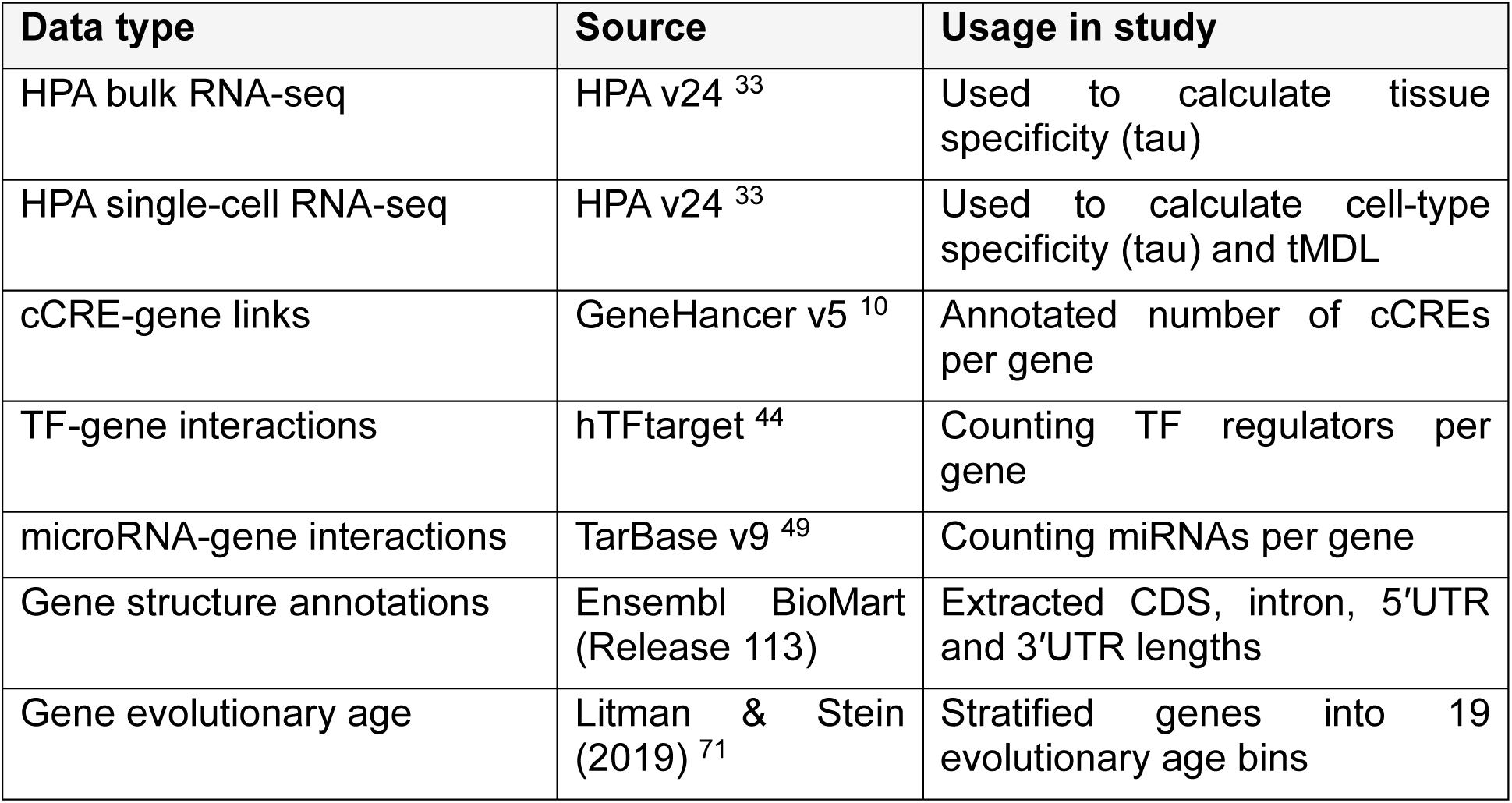

All genomic analyses were conducted using the GRCh38 (hg38) human genome assembly.

## Acknowledgements

We thank Orna Dahan, Nadav Mishol, Omer Kerner, and Zohar Yakhini for their valuable input and fruitful discussions. Y.P. is a Ben May Professorial Chair and a Kimmel Investigator. We thank the Minerva Center on Live Emulation of Evolution in the Lab and the Sharon Zuckerman Laboratory for Research in Systems Biology for grant support.

