## Supplementary Information for "An information content principle explains regulatory patterns of gene expression across human tissues"

#### Supplementary Figures

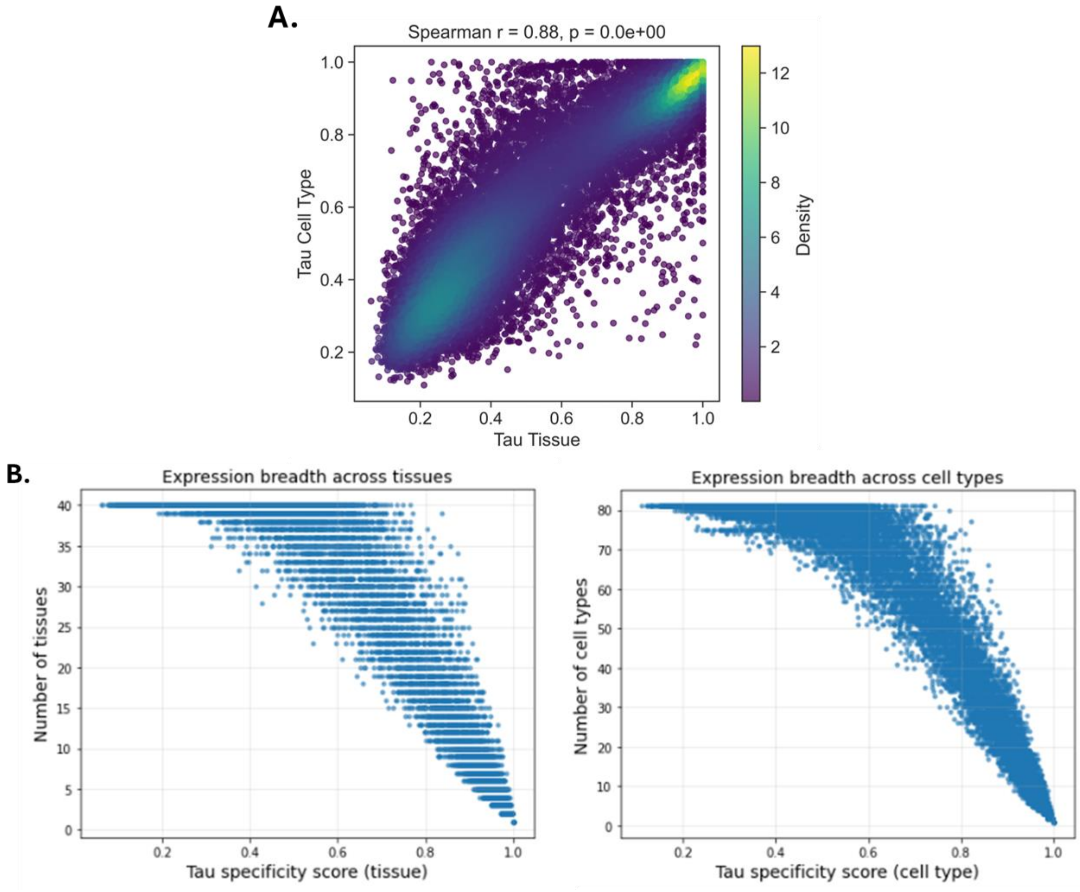


**Supplementary Figure 1. Comparison of tau values and expression breadth across bulk and single-cell datasets. (A)** Scatterplot showing the correlation between tissue-level tau (derived from bulk RNA-seq) and cell type-level tau (calculated from single-cell RNA-seq aggregated by cell type) for 18,324 human protein-coding genes. Points are colored by local density. **(B)** Scatterplots comparing tau values to expression breadth, measured as the number of tissues or cell types in which each gene is expressed. Left: tissue-level tau vs. number of tissues. Right: cell type-level tau vs. number of cell types.

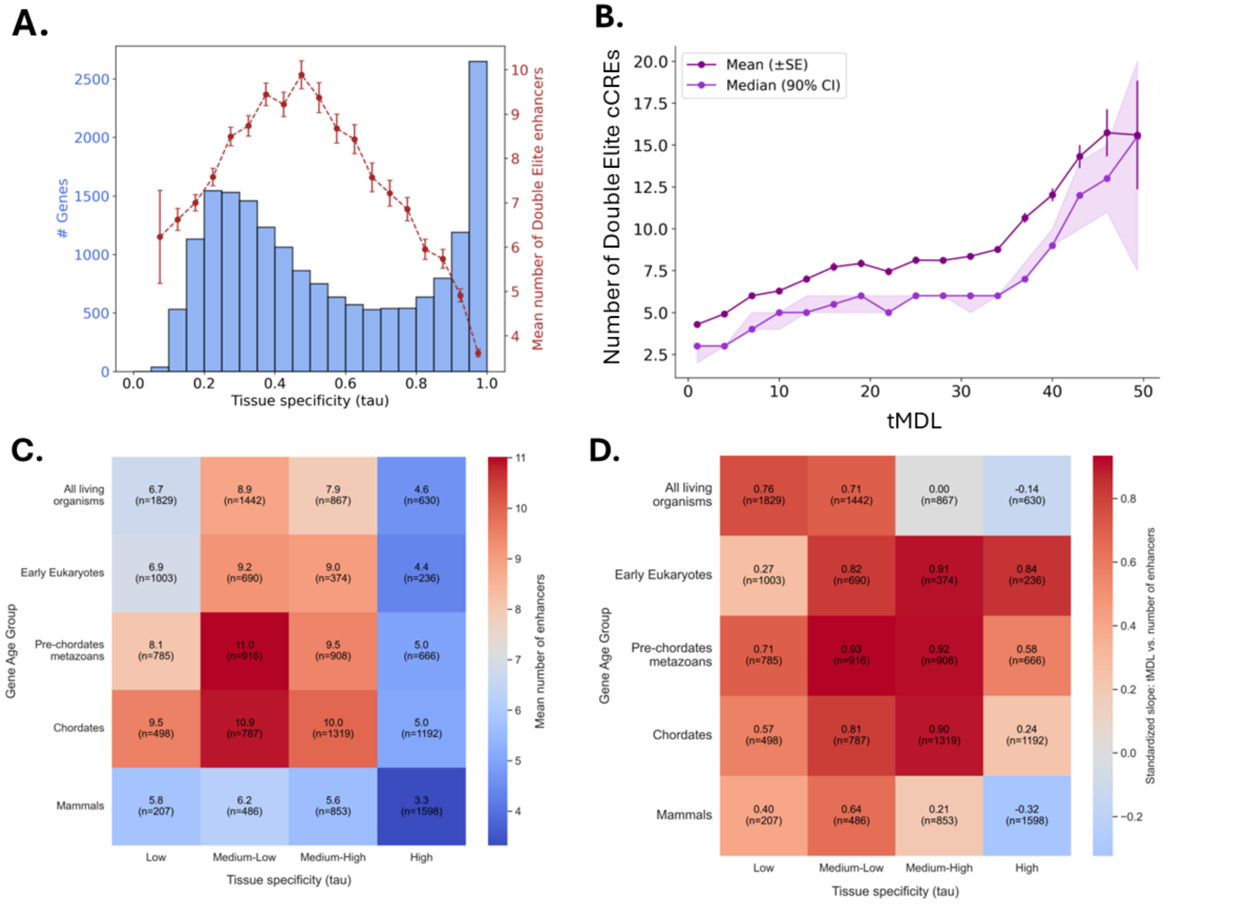


**Supplementary Figure 2. cCRE-based analyses performed on the more stringent "Double Elite" subset.** All panels replicate the analyses shown in the main text using only Double Elite cCREs, as defined by GeneHancer (see Methods), revealing patterns consistent with those observed using the full cCRE set. **(A)** Histogram of tau scores calculated from bulk RNA-seq data (blue bars, left y-axis), with overlaid line plot (red dashed line, right y-axis) indicating the mean number of linked Double Elite cCREs per gene within each tau bin. Error bars represent standard error of the mean (SEM). **(B)** Line plot showing the mean (purple) and median (violet) number of Double Elite cCREs per gene across bins of ranked tMDL values using a sliding window approach. Error bars and shaded areas represent standard error of the mean (SEM) and 90% confidence intervals (CI), respectively. **(C)** Heatmaps displaying the mean number of Double Elite CREs per gene, stratified by evolutionary age (y-axis) and tissue specificity (x-axis, binned by tau values). **(D)** Heatmaps showing the normalized slope from a sliding window analysis of Double Elite CRE number versus tMDL, stratified by evolutionary age (y-axis) and tissue specificity (x-axis, binned by tau values).

#### A. Number of bins = 3

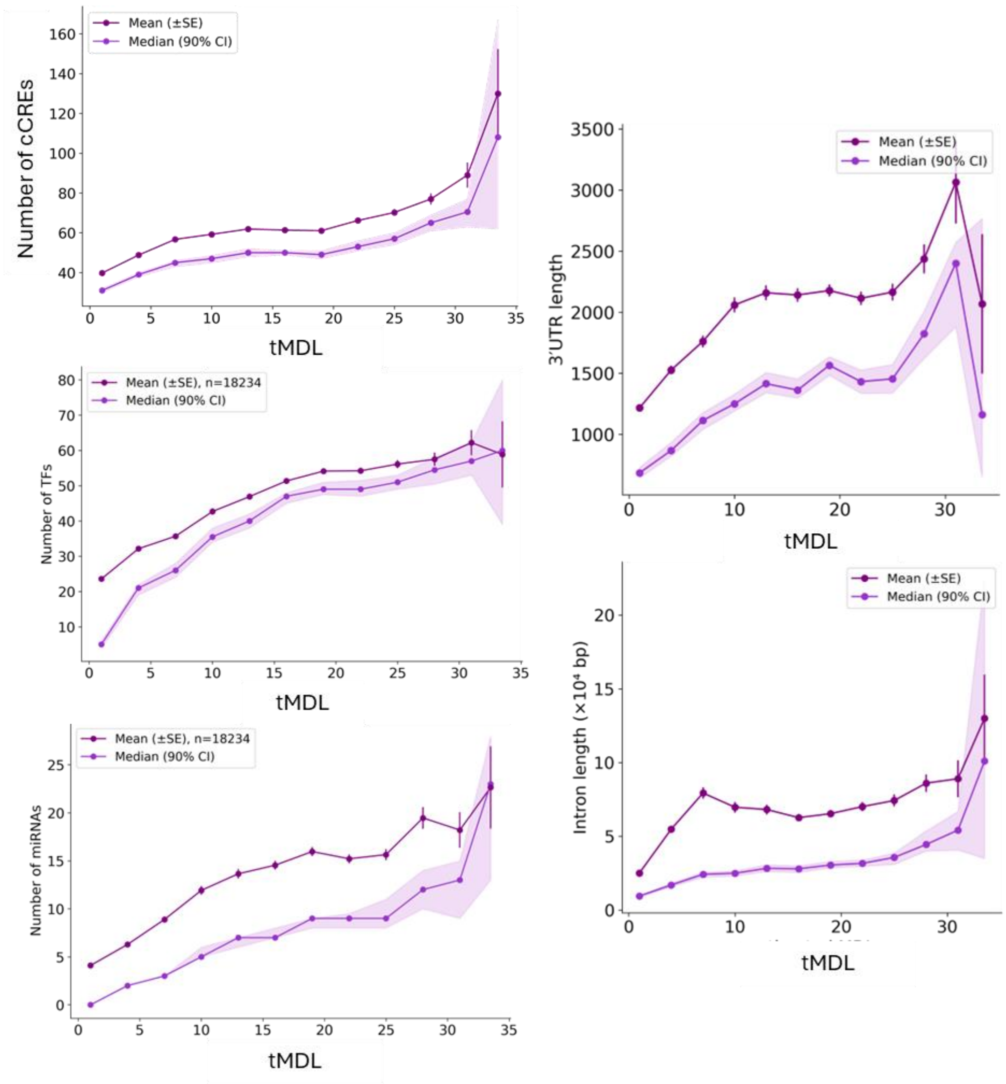

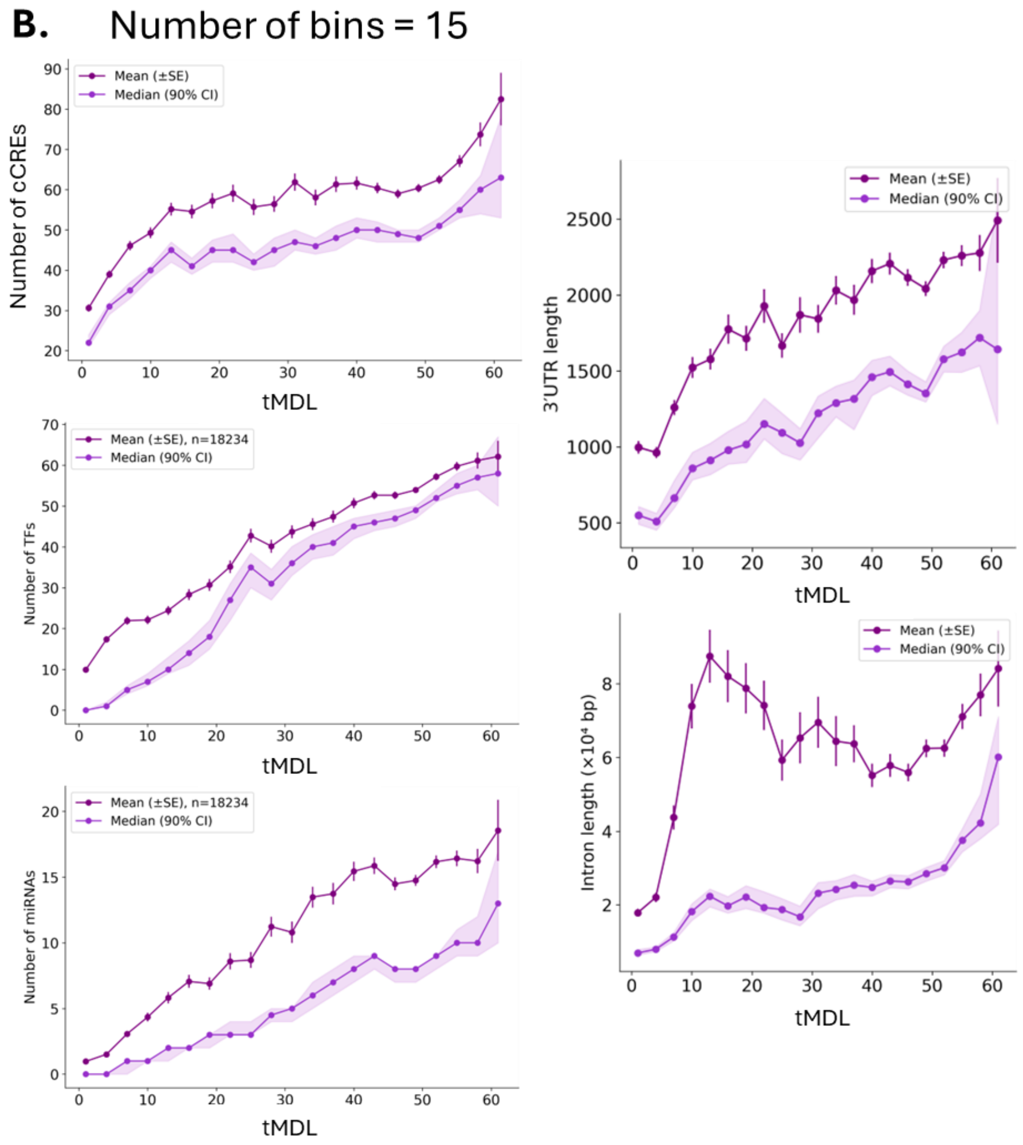


**Supplementary Figure 3. Robustness of tMDL trends to expression binning parameters.** The tMDL calculation requires discretizing continuous gene expression levels into bins, which can influence the number of inferred regulatory transitions. To test the robustness of downstream results to this choice, we repeated the tMDL-based sliding window analysis using alternative binning strategies. Line plots show the mean (purple) and median (violet) values of key regulatory features across fixed-size sliding windows of tMDL. **(A)** Results using 3 expression bins. **(B)** Results using 15 expression bins. Features shown (left to right): number of linked cCREs, number of TFs, number of miRNAs, 3′ UTR length, and intron length. In all cases, trends were overall consistent with the main analysis (six bins; see Figure 4), supporting the robustness of the tMDL framework to binning choice. Error bars and shaded areas represent standard error of the mean (SEM) and 90% confidence intervals (CI), respectively.

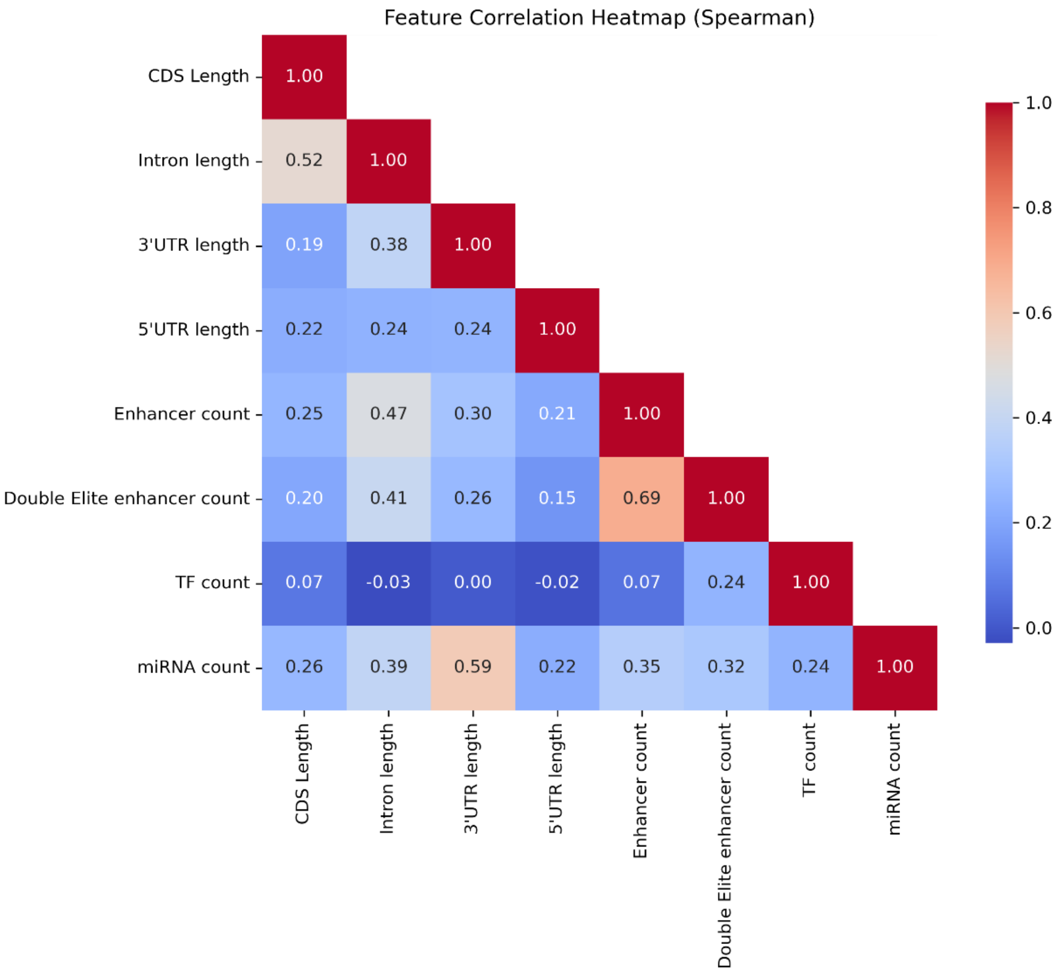


**Supplementary Figure 4. Correlation structure among gene regulatory and structural features.** Heatmap showing pairwise Spearman correlations between key gene features, including CDS length, intron length, 3′ UTR length, 5′ UTR length, number of linked cCREs (full and Double Elite subsets), and the number of TFs and miRNAs per gene. The color scale reflects the correlation coefficient (ρ).

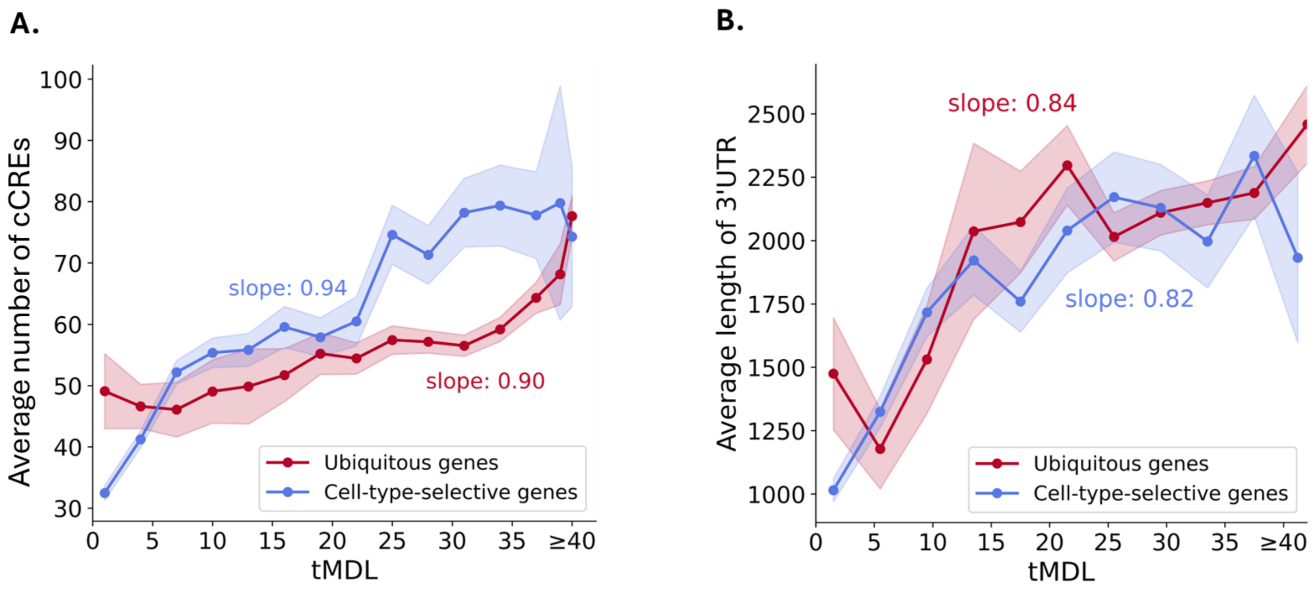


**Supplementary Figure 5. Regulatory features that scale with tMDL independently of expression breadth.** Line plots showing the relationship between tMDL and two key regulatory features, cCRE abundance and 3′ UTR length, stratified by expression breadth. Mean feature values were calculated across sliding windows of ranked tMDL, separately for ubiquitous (red) and cell-type selective (blue) genes. Slopes were obtained from linear models fit to windowed means after standardizing both variables (zero mean, unit variance) to enable comparison. Shaded areas represent 90% confidence intervals (CI). To avoid sparse windows, all genes with tMDL ≥ 40 were grouped into a final bin. Both cCRE count and 3′ UTR length increased with tMDL in each regime, highlighting their universal contribution to regulatory complexity across diverse expression strategies.

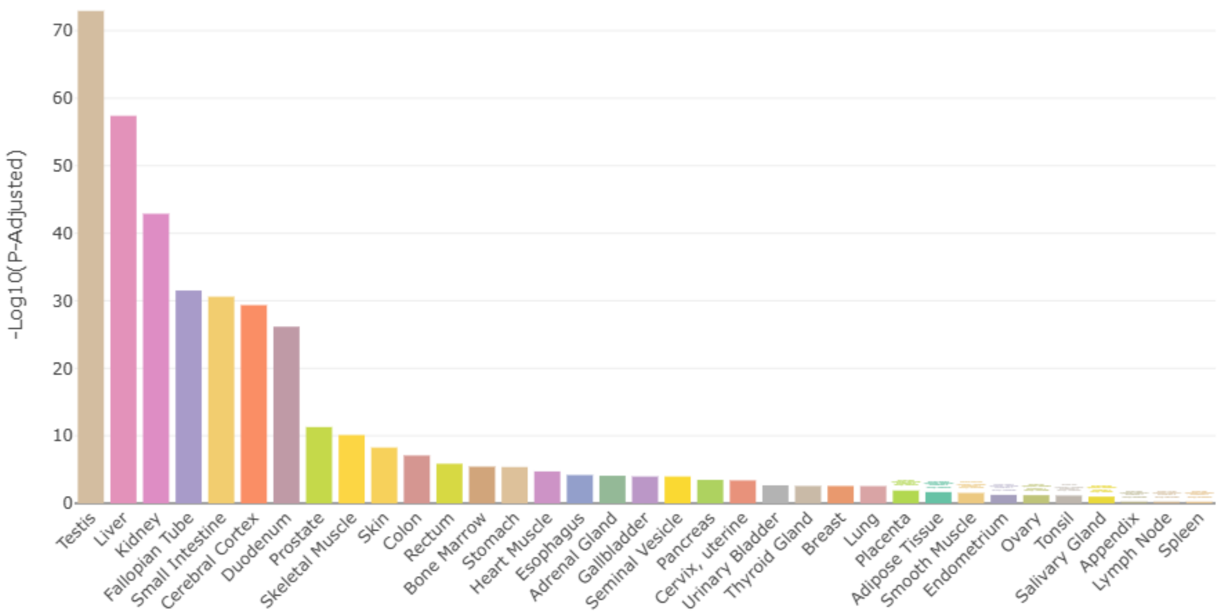


**Supplementary Figure 6. Tissue enrichment of ancient tissue-specific genes.** Bar plot showing tissue enrichment analysis results for ancient genes with tissue-specific expression patterns. Enrichment was assessed using TissueEnrich, which tests for overrepresentation of input genes among tissue-enriched gene sets defined in the Human Protein Atlas. The three top enriched tissues are testis, liver, and kidney.

A.

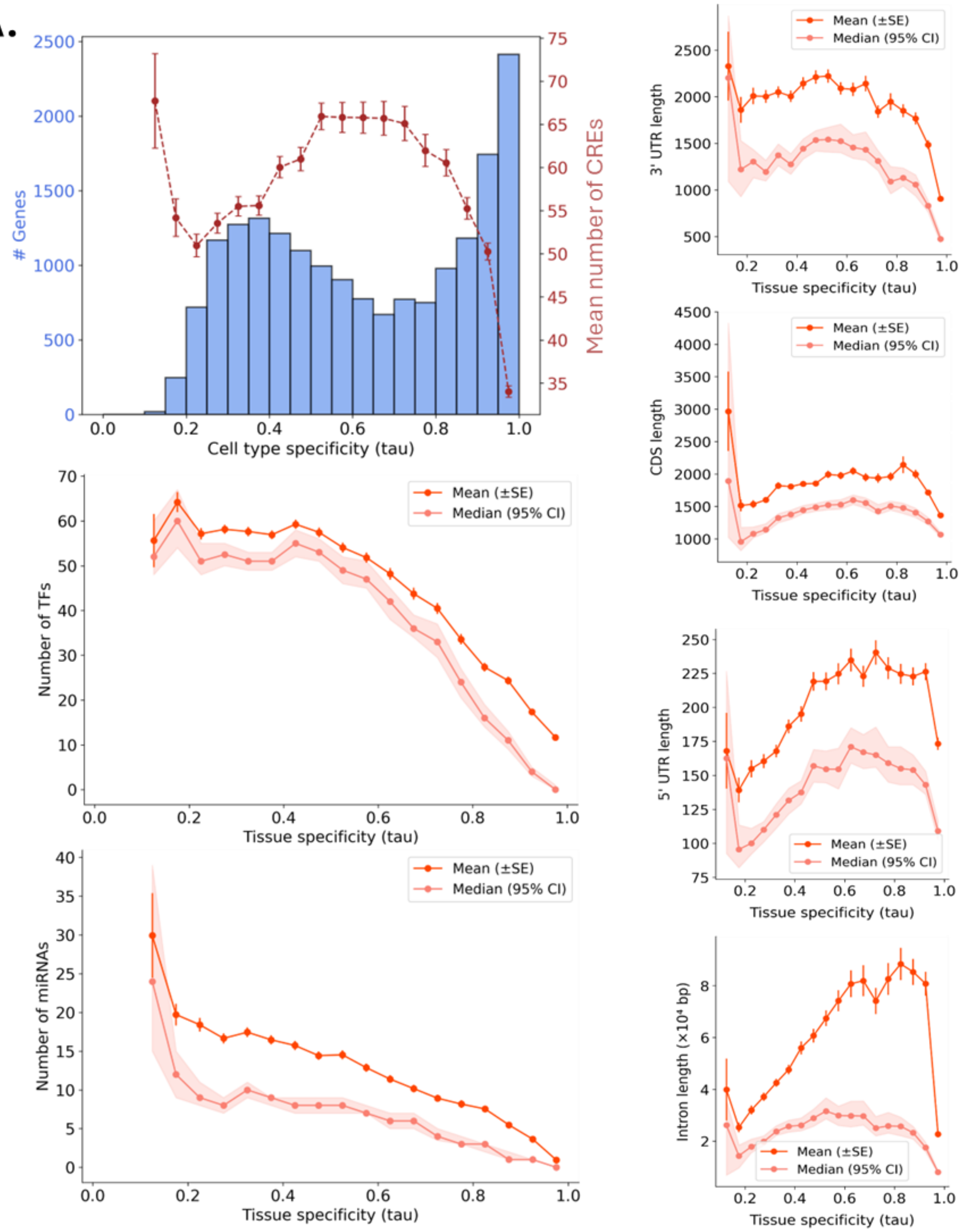

### Genes
Mean number of CREs
Cell type specificity (tau)
Number of TFs
Number of miRNAs
Tissue specificity (tau)
Mean (±SE)
Median (95% CI)
3' UTR length
CDS length
5' UTR length
Intron length (×10⁴ bp)

B.

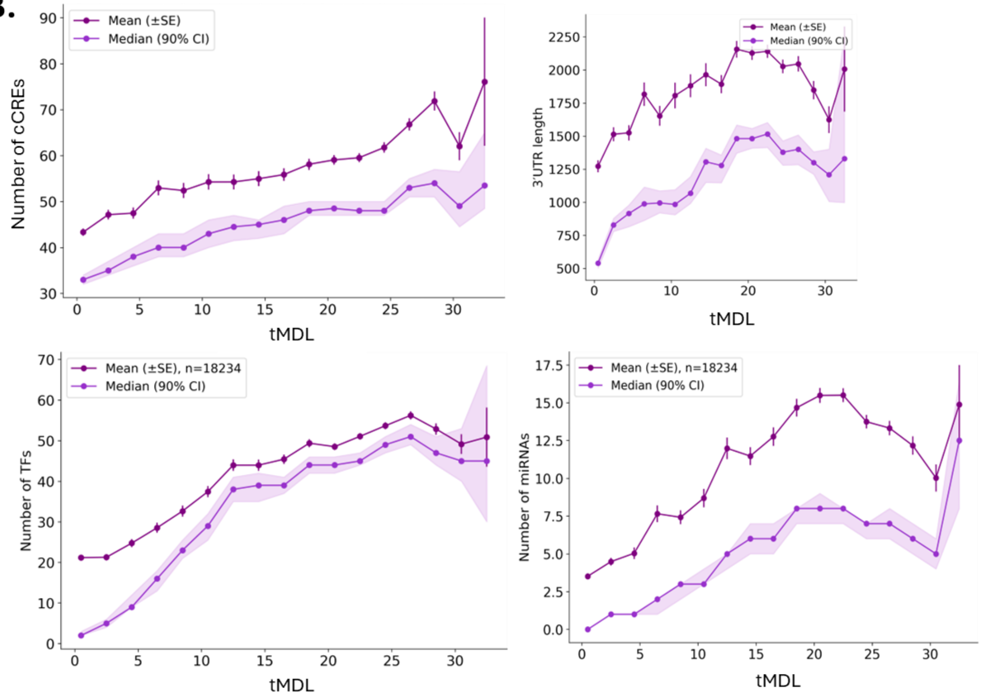


**Supplementary Figure 7. Robustness of regulatory and structural feature trends across tissue-level and cell type-level definitions of tau and tMDL. (A)** Regulatory and structural feature trends across tau calculated from single-cell RNA-seq data aggregated by cell type, showing overall agreement with results using tissue-level tau. The first panel shows a histogram of tau values (blue bars, left y-axis) with an overlaid line plot (red dashed line, right y-axis) indicating the mean number of linked cCREs per gene within each tau bin. Subsequent panels show the mean (purple) and median (violet) values of TF count, miRNA count, 3′ UTR length, CDS length, 5′ UTR length, and intron length across sliding windows of tau. **(B)** Regulatory feature trends across tMDL calculated from bulk RNA-seq tissue data, showing overall agreement with the main analyses based on cell-type–derived tMDL. Line plots display the mean and median number of linked cCREs, TFs, miRNAs, and 3′ UTR length across sliding windows of tMDL. No consistent signal was observed for intron length in this tissue-level analysis. Error bars and shaded areas represent standard error of the mean (SEM) and 90% confidence intervals (CI), respectively.
